# A “spindle and thread”-mechanism unblocks translation of N-terminally disordered proteins

**DOI:** 10.1101/2021.02.22.432221

**Authors:** Margit Kaldmäe, Thibault Vosselman, Xueying Zhong, Dilraj Lama, Gefei Chen, Mihkel Saluri, Nina Kronqvist, Jia Wei Siau, Aik Seng Ng, Farid J. Ghadessy, Pierre Sabatier, Borivoj Vojtesek, Médoune Sarr, Cagla Sahin, Nicklas Österlund, Leopold L. Ilag, Venla A. Väänänen, Saikiran Sedimbi, Roman A. Zubarev, Lennart Nilsson, Philip J. B. Koeck, Anna Rising, Nicolas Fritz, Jan Johansson, David P. Lane, Michael Landreh

## Abstract

Protein disorder is a major hurdle for structural biology. A prominent example is the tumour suppressor p53, whose low expression levels and poor conformational stability due to a high degree of disorder pose major challenges to the development of cancer therapeutics. Here, we address these issues by fusing p53 to an engineered spider silk domain termed NT^*^. The chimeric protein displays highly efficient translation *in vitro* and in *E. coli* and is fully active in human cancer cells. The transmission electron microscopy structure and native mass spectrometry reveal that the full-length p53 fusion protein adopts a compact conformation. Molecular dynamics simulations show that the disordered transactivation domain of p53 is wound around the NT^*^ domain via a series of folding events, resulting in a globular structure. We find that expression of B-Raf, another partially disordered cancer target, is similarly enhanced by fusion to NT^*^. In summary, we demonstrate how inducing co-translational folding via a molecular “spindle and thread” mechanism can overcome poor translation efficiency of partially disordered proteins.

## Introduction

Proteins with intrinsically disordered domains (IDDs) that do not adopt a well-defined three-dimensional structure are involved in a variety of essential cellular processes, but are significantly enriched among cancer-related proteins (Santofimia-Castaño et al., 2020; Uversky et al., 2008). The ability of IDDs to participate in promiscuous interactions ranging from transcription factor recruitment to cell signalling make them attractive targets for therapeutic approaches, yet complicates structural biology and drug design efforts (Tsafou et al., 2018). Perhaps the most prominent example is the tumor suppressor p53, which plays a central role in the control of cell proliferation (Lane, 1992). Mutations that inactivate p53 can be found in up to 60% of all human cancers, and consequently, restoring its function is an attractive therapeutic strategy (Brown et al., 2013; Bykov et al., 2018; Chang et al., 2013; Joerger and Fersht, 2016; Zawacka-Pankau and Selivanova, 2015). Its N-terminal region, composed of transactivation domains 1 and 2 (TAD1 and TAD2, residues 1–40 and 41–60) and a proline-rich domain (PRD, residues 61–92), is intrinsically disordered (Wells et al., 2008). N-terminal disorder is a negative regulator of protein half-life in human cells and can be considered a hallmark of proteins that are subject to rapid turnover, in agreement with the short life cycle of p53 in healthy cells (van der Lee et al., 2014). As a result, the basal expression levels of p53 in human cells are low, providing little evolutionary pressure to adopt a stable structure (Vecchi et al., 2020). Consequently, p53 displays high aggregation propensity *in vitro* and can spontaneously assemble into amyloid-like fibrils (Silva et al., 2014).

These observations raise the question whether the N-terminal disorder could be targeted to produce a stably folded p53 variant. To find out how evolution has addressed this issue, we turned to a protein system that contains highly disordered, aggregation-prone segments yet is expressed at extremely high levels. Major ampullate spidroins, the proteins that form spider dragline silk, are expressed in the silk gland and stored in soluble form in the sac at concentrations exceeding 40% *w/v*. They contain a highly conserved N-terminal domain (NT) composed of 130 residues, followed by several hundred repeats of poly-alanine and glycine-rich stretches (Fig. 1 a), and a regulatory C-terminal domain (Babb et al., 2017). The tightly folded, α-helical NT mediates spidroin assembly through pH-dependent dimerization (Kronqvist et al., 2014; Landreh et al., 2010). We recently showed that a non-dimerizing mutant (NT^*^) enables efficient production of aggregation-prone peptides by increasing expression while suppressing self-assembly (Abelein et al., 2020; Kronqvist et al., 2017; Sarr et al., 2018). We therefore speculated that the specific properties of NT^*^ could be utilized to modulate the N-terminal disorder of p53 and other partially disordered proteins to obtain previously inaccessible structural and functional insights.

**Figure 1.**
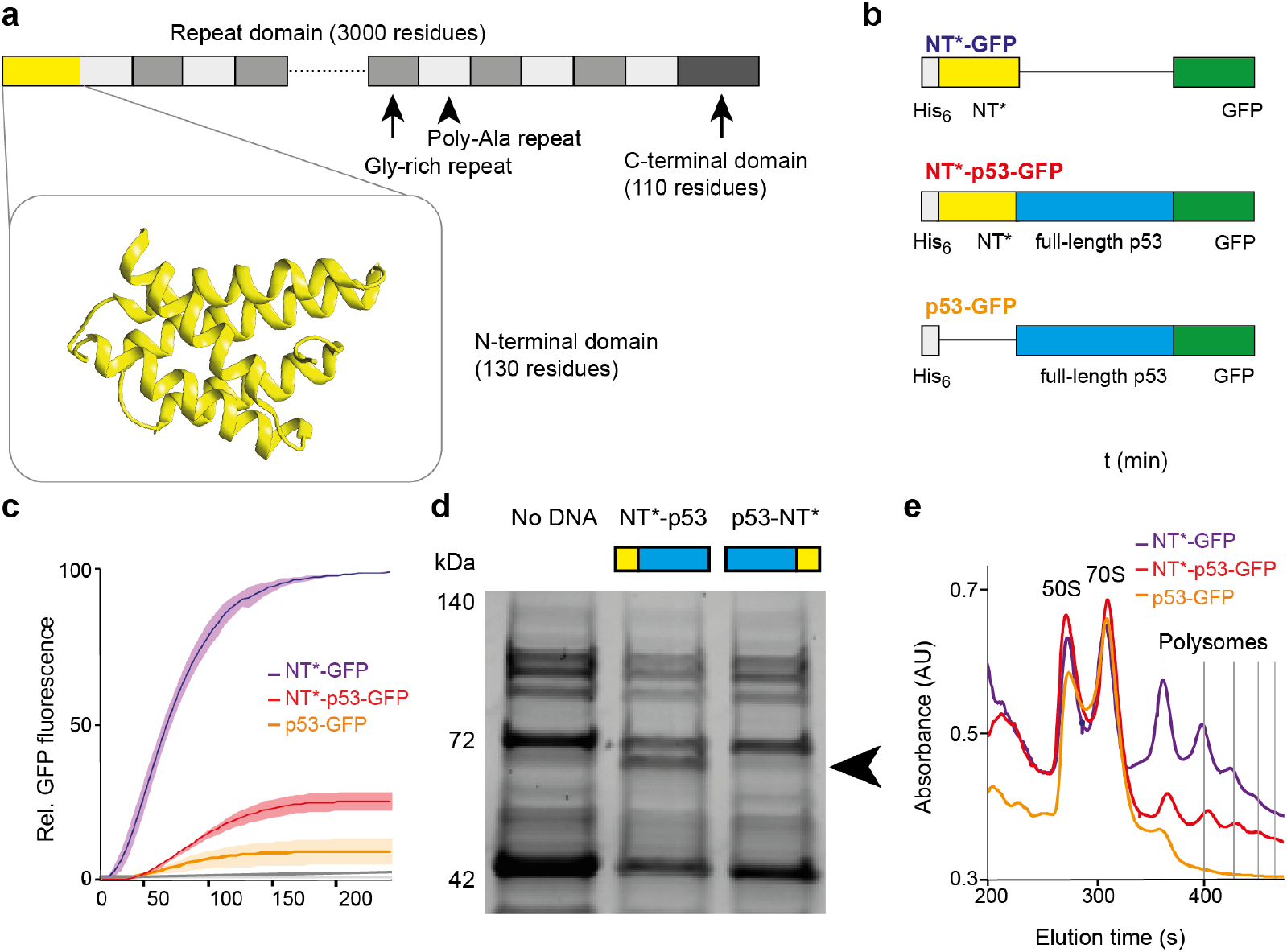
NT^*^ increases the translation efficiency of p53. (a) Architecture of major ampullate spidroin (MaSp) proteins. The NT domain (yellow, PDB ID 2LPJ) is followed by several hundred low-complexity repeats of alternating poly-Ala and Gly/Gln-rich (MaSp1) or Pro/Gly-rich (MaSp2) segments, and a small regulatory C-terminal domain. (b) Constructs used for IVT. (c) IVT of the GFP-tagged NT^*^- and p53-constructs. Curves show the mean GFP fluorescence intensity as a function of time. Shaded areas indicate standard deviations (n=3). (d) SDS-PAGE analysis of IVT reactions expressing NT^*^-p53 and p53-NT^*^ shows that only the N-terminally tagged p53 is expressed at significant amounts. Arrows indicate the approximate location of the fusion protein band. (e) Polysome profiles of the IVT reactions in (c) after 100 min. The positions of the 50S and 70S ribosomes, as well as polysomes composed of up to six ribosomes are indicated.

## Results

### NT^*^ increases translation efficiency of p53

Translation represents a basal level at which protein expression can be regulated. We therefore first investigated the effect of NT^*^ on p53 translation. We designed DNA constructs that encompassed either an NT^*^ domain, full-length p53, or NT^*^ fused to the N-terminus of p53 (Fig. 1 b, Fig. S1 a). All constructs contained a C-terminal GFP tag (Table S1). We then performed *in vitro* transcription and translation (IVT) of each construct using a reconstituted bacterial protein synthesis system and monitored GFP fluorescence intensity over time. The approach is thus independent of biological limitations such as protein toxicity and depletion of rare tRNAs, and allows an unbiased assessment of translation efficiency (Afonina et al., 2015). As expected, IVT of NT^*^-GFP showed a robust increase in GFP fluorescence with a lag phase of less than 10 min. Translation of p53-GFP, on the other hand, had a longer lag phase of 45 min and a maximum fluorescence intensity 90% below that of NT^*^-GFP. IVT of NT^*^-p53-GFP exhibited a 30 min lag phase and reached a maximum fluorescence intensity of approximately 25% relative to NT^*^-GFP (Fig. 1 c). To probe the importance of the location of the NT^*^ tag, we compared translation levels of p53 with an N-terminal or a C-terminal NT^*^ tag and no GFP. Strikingly, only the N-terminally tagged protein could be detected (Fig. 1 d). These findings suggest that an N-terminal NT^*^ increases translation of p53 *in vitro*.

Next, we asked whether the differences in GFP fluorescence between p53-GFP and NT^*^-p53-GFP could be due to differences in ribosome activity. We therefore performed polysome profiling of the IVT reactions after 100 mins, when all three constructs showed a linear increase in fluorescence. For the NT^*^-GFP and NT^*^-p53-GFP, we detected up to 6 translating ribosomes per RNA, whereas the polysome profile of the p53-GFP RNA showed a maximum of two bound ribosomes per RNA, indicating that translation of p53 is either slowed down or stalled in the absence of NT^*^ (Fig. 1 e). Since the IVT system employed here is derived from the protein synthesis machinery of *E. coli*, we also compared the expression of p53 and NT^*^-p53 without GFP tags in the *E. coli* expression strain BL21(DE3). In line with the IVT results, we found that NT^*^-p53, but not p53 alone, is expressed highly efficiently, yielding up to 2 g of crude NT^*^-p53 inclusion bodies per 1 L of bacterial culture, as opposed to low mg amounts for untagged p53 (Fig. S1 b). We then tested expression of the disordered TAD domains (residues 1 – 60) of p53 with and without NT^*^. Similar to its effect on the expression of full-length p53, NT^*^ increased the protein yield of NT^*^-TAD1-TAD2 approximately five times compared to TAD1-TAD2 alone (Fig. S1 c).

### NT^*^-p53 forms a native tetramer

Having established that an N-terminal NT^*^ domain increases p53 expression, we set out to determine how it affects the structure of p53. For this purpose, we could take advantage of the high expression levels afforded in *E. coli* to purify recombinant NT^*^-p53 for structure analysis. We then established a purification protocol which yielded >90% pure NT^*^-p53 through preparation and refolding of inclusion bodies (Fig. S2 a, b). The formation of the native tetramer was confirmed through glutaraldehyde crosslinking and Western blot analysis. Exclusively monomeric protein was detected if the protein was denatured in the absence of glutaraldehyde, while the crosslinked protein remained tetrameric (Fig. S3 a). To confirm the correct structure of the refolded protein, we performed antibody pull-down experiments using DO-1, which recognizes the N-terminal MDM2 binding site (Vojtěšek et al., 1992), PAb 1620, which recognizes the folded core domain (Ball et al., 1984), PAb 421, which recognizes the C-terminal domain (Harlow et al., 1981), and PAb 240, which recognizes a cryptic epitope in partially unfolded p53 (Gannon et al., 1990). Briefly, antibodies immobilized on magnetic beads were incubated with recombinant NT^*^-p53 and the presence of p53 was confirmed by Western blot analysis. In all cases, we could detect p53, indicating the specific epitope of each antibody is present in the refolded NT^*^-p53 fusion protein (Figure 2 a). These findings suggest that the NT^*^ domain does not alter the conformation or oligomerization of the p53 moiety.

**Figure 2.**
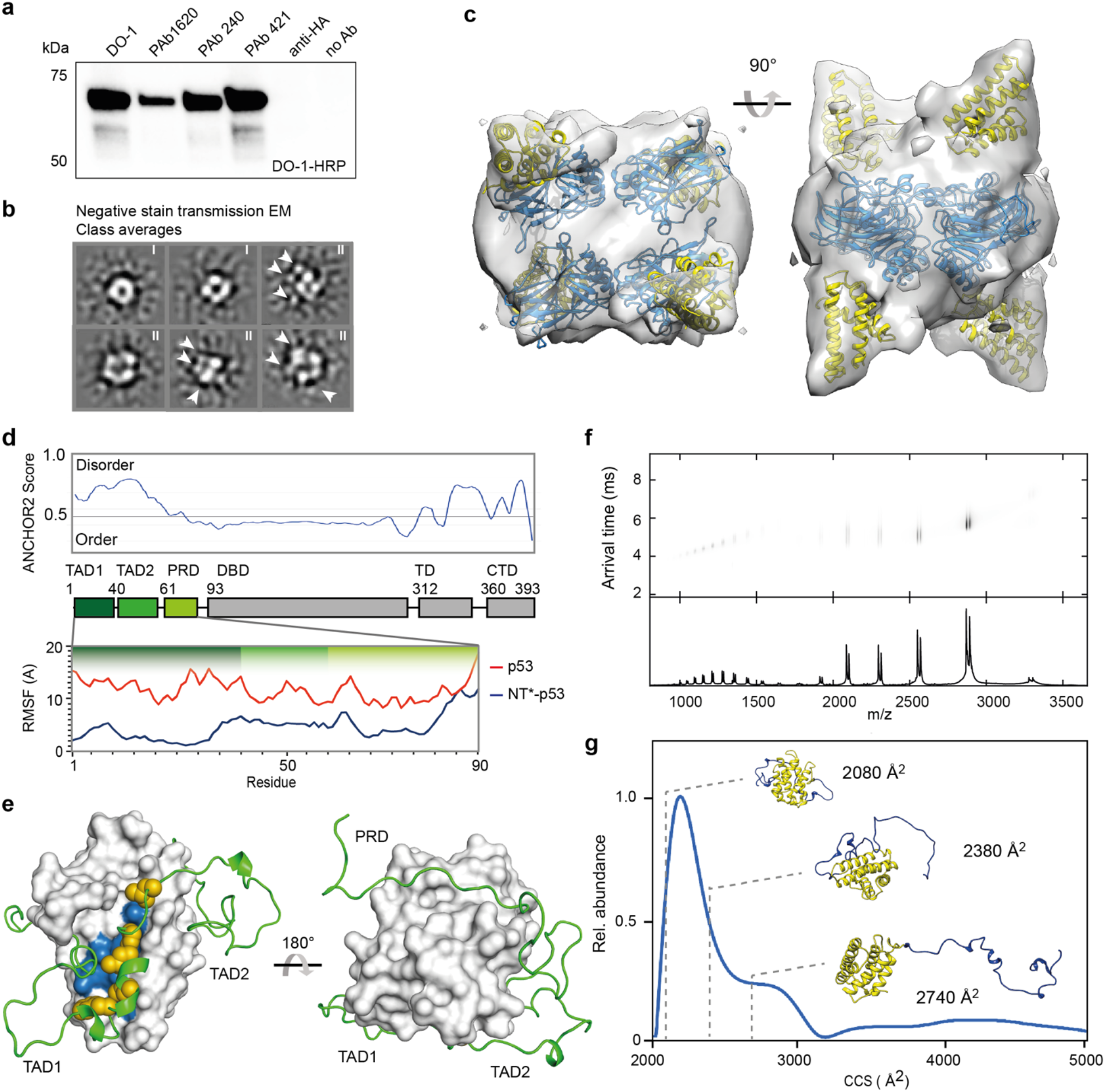
The conformation of the NT^*^-p53 tetramer with compact TADs. (a) Western blot of refolded NT^*^-p53 after immunoprecipitation with an antibody recognizing the MDM2 binding site (DO-1), as well as conformation-specific antibodies to the folded and unfolded core domain (PAb1620, and PAb240), and the C-terminal domain (PAb421) show that NT^*^-p53 displays the epitopes associated with folded p53. (b) TEM class averages of NT^*^-p53 show a ring-like structure with four individual domains (I), with can be surrounded by one to three smaller densities (II). (c) 3D reconstruction of NT^*^-p53 shows a central core composed of four DNA binding domains (blue, PDB ID 3KMD) with four extra densities corresponding to NT^*^ domains (yellow, PDB ID 2LPJ). (d) Overall architecture and disorder of p53. The three disordered N-terminal parts, TAD1, TAD2, and PRD, are indicated in dark, medium, and light green, respectively. Insert: RMSFs of the p53 N-terminus with or without an N-terminal NT^*^ domain over the course of a 1 μs MD simulation. (e) End-point representative structure of the N-terminal region of the NT^*^-p53 fusion construct from all-atom simulations shows wrapping of the domains around NT^*^. NT^*^ is shown as surface representation with the hydrophobic patch formed by residues M41, I44, M48, I52, L55, L66, A68, L69, A72, F73 and M77 coloured in blue. The N-terminal domain of p53 is shown as ribbon rendering. Hydrophobic residues (F19, L22, L26, P27 and L32) in the TAD1 binding site are shown as yellow spheres. (f) IM-MS analysis of recombinant NT^*^-TAD1-TAD2 protein reveals mostly lowly charged, compact conformations. (g) Plotting the CCS distributions of NT^*^-TAD1-TAD2 reveals that most of the protein has a CCS centred around 2100 Å^2^, in good agreement with the compact conformation predicted by MD simulations. A minor population exhibits a more extended conformation, indicating TAD domain unfolding. The NT moiety is shown in yellow and the TAD domains are shown in blue.

Due to its partially disordered nature, no high-resolution structure of full-length p53 has been solved to date. However, transmission electron microscopy (TEM) studies have revealed that in the absence of DNA, p53 adopts a ring-like structure with the four core domains arranged around a central cavity (Pham et al., 2012). Since the NT^*^ domain with 130 residues is only moderately smaller than the 312 residue-core domain of p53, we investigated the orientations of the NT^*^ tags in the NT^*^-p53 tetramer using TEM. Analysis of the class averages calculated from 6519 manually picked particles showed a ring-shaped four-domain architecture as previously observed in TEM of wt p53 (Fig. 2 b, Fig. S3 b). However, in most classes, we could detect additional, smaller densities near the central ring. Reconstruction of the NT^*^-p53 tetramer based on the densities observed by TEM yielded a compact model of the NT^*^-p53 complex with a resolution of 13 Å (Fig. 2 c, Fig. S3 c, d). In this model, each DNA binding domain of the p53 tetramer is closely associated with an NT^*^ domain. The NT^*^ domains on diametrically opposed domains are located on the same side of the core of the complex, with two above and two NT^*^ domains below the ring. We therefore concluded that the disordered N-terminal region of p53 is not extended in these complexes.

### NT^*^ induces compaction of the TADs

To obtain more detailed insights into the effect of NT^*^ on the disordered p53 N-terminus, we performed all-atom molecular dynamics (MD) simulations of the first 90 residues of p53, covering the disordered region with both TADs and the PRD, with or without the NT^*^ domain. Comparisons of root mean square fluctuations (RMSF) of p53 N-terminal regions between the two constructs as a measure of the degree of conformational flexibility show that the NT^*^-linked variants exhibit significantly lower RMSFs (Fig. 2 d). The shift in RMSF is due to the p53 polypeptide linked with NT^*^ wrapping around the protein; except for the residues towards the terminal end of PRD which remained largely unbound over the course of the simulation (Fig. 2 d). Thus, the presence of NT^*^ significantly reduces disorder from the intrinsically disordered N-terminus of p53. Folding of the p53 TADs and PRD onto NT^*^ are likely driven by hydrophobic collapse, as the hydrophobic residues present along the TAD1 domain are observed to interact with a predominantly hydrophobic patch on the surface of NT^*^ that is part of the dimerization interface in the wt NT domain (Figure 2 e). The hydrophobic nature of the association between TAD1 and NT^*^ was consistently observed in multiple simulations (Fig. S4 a - e). However, the exact orientations of the bound TAD1 domain differed between these simulations, indicating low specificity.

To confirm the interactions between NT^*^ and the TADs, we expressed and purified TAD1 and TAD2 with and without an N-terminal NT^*^ domain and used ion mobility mass spectrometry (IM-MS) to assess the conformation of the fusion protein. By separating protein conformers in a gas-filled flight tube, we are able to measure their collision cross sections (CCS) and relative abundances. IM-MS analysis of NT^*^-TAD1-TAD2 shows a major population with a CCS of 2100 Å^2^, in good agreement with the 2080 Å^2^ predicted from the MD simulations. We also observed a minor population around 2700 Å^2^, which corresponds to NT^*^ with an extended TAD1-TAD2 moiety, as well as traces of higher unfolding (Fig. 2 f, g). Together, the insights from TEM and MD simulations indicate that NT^*^ remains folded in the p53 fusion protein and induces a compact conformation of the chimeric construct in which the p53 N-terminal transactivation region wraps around NT^*^.

### NT^*^-p53 is active in human cancer cell lines

Based on our observation that NT^*^ induces a compactly folded state of p53, we then asked whether the fusion protein would retain its functionality in human cancer cells. To address these questions, we generated vectors for mammalian cell expression of p53, NT^*^-p53, and p53 with the Arg273His mutation (p53R273H) that inactivates p53 by impairing DNA binding, under the CMV promotor. All constructs were transfected into p53 null non-small cell lung carcinoma (H1299) cells, and the subcellular localization of the proteins monitored by immunohistochemistry of transfected H1299 cells with DO-1. We found that p53 was located predominantly in the nucleus, the expected localization for the wt protein. For NT^*^-p53, we observed cells with nuclear staining, as well as a small population of cells with nuclear and cytosolic NT^*^-p53 (Fig. 3 a). The even staining and the absence of speckles indicate that the NT^*^-p53 fusion protein does not aggregate in the cell and can be imported into the nucleus in the same manner as wt p53.

**Figure 3.**
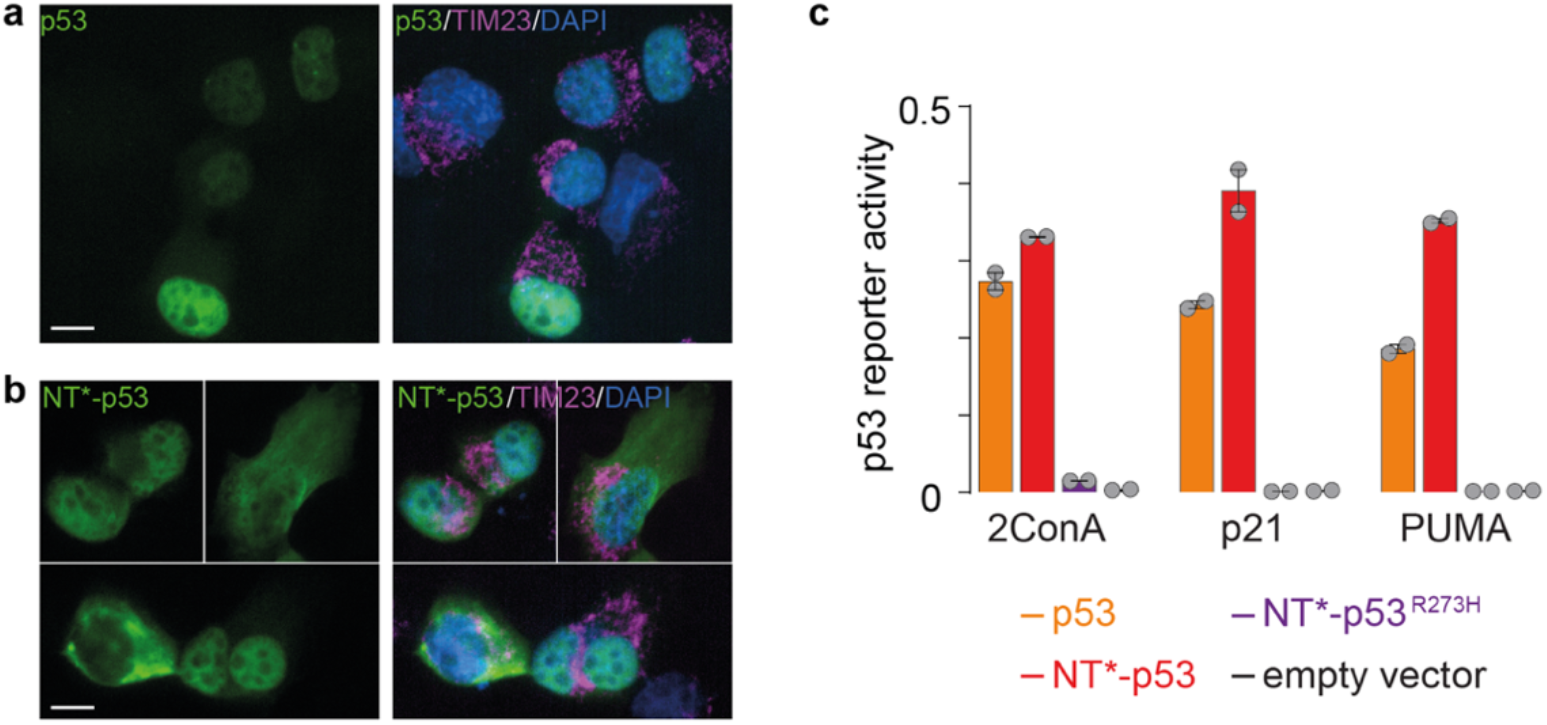
NT^*^-p53 is active in H1299 cells. (a, b) Representative confocal immuno-cytochemistry pictures of H1299 cells 8 hrs after transfection with the indicated constructs and stained with anti-p53 (DO1, green) and anti-TIMM23 (magenta, indicates mitochondria) antibodies. DAPI staining was used to label the nuclei. Scale bars, 10 mm. In (b), a montage of three pictures was used to accurately represent the data. The data is representative of 2-3 independent experiments. (c) Bar chart of the mean ± standard error to the mean relative p53 reporter gene activity in H1299 cells expressing p53, NT^*^-p53, p53^R273H^, or the empty vector (n=2). Grey dots represent independent experiments.

To assess whether NT^*^-p53 specifically induces targets related to cell cycle arrest and apoptosis, we monitored the activity of the different p53 fusion proteins in H1299 cell lines carrying luciferase reporters for tandem copies of a synthetic p53 consensus response element (2ConA), apoptosis mediator PUMA, and cell cycle arrest factor p21WAF1 (Carbone et al., 1992). Here, p53 and NT^*^-p53 induced comparable levels of luciferase driven by the high affinity 2ConA response element. Interestingly, NT^*^-p53 expression resulted in a 1.3-fold higher induction of p21WAF1 and 1.5-fold higher induction of PUMA reporters compared to p53 without NT^*^ (Fig. 3 b). The R273H mutant, on the other hand, was inactive. However, we did not detect higher NT^*^-p53 expression by Western blot, suggesting that the difference in activity is not dependent on protein levels (Figure S5). Taken together, these findings suggest that the compact TAD conformation induced by the NT^*^ domain does not impair the transcription factor activity of p53 in human cells.

### NT^*^ promotes translation of disordered B-Raf but not folded K-Ras

Considering that NT^*^ increases expression of p53 *in vitro* and in *E. coli*, while at the same time inducing a compact conformation of the TAD, we speculated about a causal relationship between both observations. Specifically, we asked whether the interactions with NT^*^ could boost translation of other N-terminally disordered proteins. To test this hypothesis, we selected two proto-oncogenes, the serine/threonine-protein kinase B-Raf and the GTPase K-Ras 4b. Like p53, B-Raf contains a disordered N-terminal region followed by a folded domain. K-Ras, on the other hand, has a stably folded N-terminus and a short, disordered tail (Fig. 4 a). We generated vectors for both proteins with and without an N-terminal NT^*^ domain and expressed both in *E. coli* under same conditions as NT^*^-p53. Strikingly, NT^*^-B-Raf was readily expressed at high amounts, while the untagged protein could not be detected. K-Ras, in the other hand, showed high expression levels independently of being fused to an NT^*^domain (Fig. 4 b).

**Figure 4.**
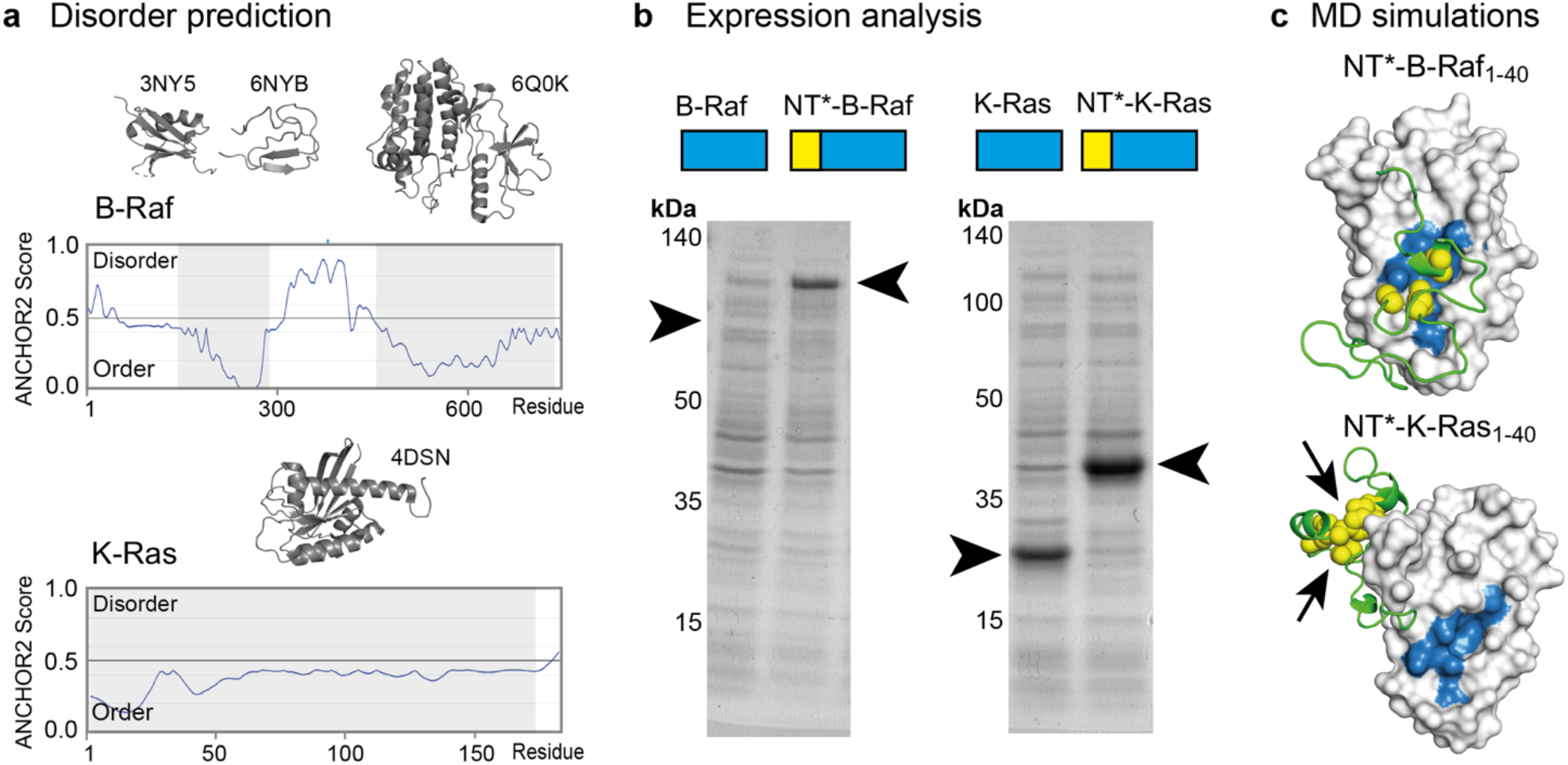
Application of the NT^*^ strategy to B-Raf and K-Ras. (a) Disorder predictions show a disordered N-terminus for B-Raf, whereas K-Ras has a folded N-terminal domain. Folded regions and the corresponding high-resolution structures and PDB IDs are shown in cartoon representation coloured in grey. (b) SDS-PAGE analysis of lysates from *E. coli* expressing wt and NT^*^-tagged B-Raf or K-Ras shows that B-Raf requires fusion to an NT^*^ domain, whereas K-Ras is readily detected without. (c) End state representative structures from MD simulations of NT^*^ fused to the first 40 residues of B-Raf (top) or K-Ras (bottom). The disordered B-Raf N-terminus wraps around the NT^*^ domain which is shown in surface representation and interacts with hydrophobic surface patch, depicted with the same set of residues as in Figure 4 coloured in blue. The K-Ras N-terminus does not bind to the patch and adopts instead a molten globular conformation with hydrophobic residues at its core (arrows). Hydrophobic residues on B-Raf (L3, A16, L17) and K-Ras (V8, V9, I21, I24, Y32, Y40) are rendered as yellow spheres.

We then asked whether the N-termini of both proteins differed in their propensity to interact with NT^*^. Analogous to our approach to the NT^*^-p53, we performed multi-copy MD simulations of the first 40 residues of either B-Raf or K-Ras, starting from an extended state and fused to an N-terminal NT^*^ domain, and followed the conformational ensembles occupied by the fusion protein. The N-terminal of B-Raf adopted a wrapped configuration highly similar to that of the p53 TAD1, mediated by multiple interactions with the hydrophobic surface patch on the NT^*^ domain (Fig. 4 c and Fig. S6). K-Ras, however, did not form extensive contacts with the hydrophobic patch on NT^*^, assuming instead a separate, molten globular structure around a hydrophobic core. Considering the observations for both protein systems, we find that NT^*^ increases expression of the partially disordered B-Raf while inducing a compact conformation of the flexible N-terminus, as seen for p53. The expression of K-Ras, which has a high propensity to form a well-folded globular structure on its own due to the high number of hydrophobic residues and low proline content, is not affected by NT^*^ in a similar way. These observations lend support to the hypothesis that the NT^*^-mediated increase in translation is related to a reduction in N-terminal disorder in the target proteins.

## Discussion

IDDs are a common feature of cancer-related proteins but often associated with a short halflife in the cell and reduced stability *in vitro*. In case of p53, these features have rendered the full-length protein refractory to structural investigations. Here, we report that a spider silk domain boosts translation of p53 as well as the similarly disordered B-Raf and induces compact conformations in their intrinsically disordered N-terminal domains, overcoming both limitations.

To unravel the connection between NT^*^, translation, and N-terminal disorder, we first considered the effect of NT^*^ on p53 translation. Using an *in vitro* translation and ribosome profiling strategy designed to probe the efficiency of translation, we find that the p53 sequence is not translated with high efficiency, in line with the fact that disordered proteins with a high proline content are poorly translated (Pavlov et al., 2009). For example, p53 translation *in vivo* depends on the elongation factor eIF5A, which suppresses ribosomal pausing (Gutierrez et al., 2013; Li et al., 2004). Co-translational folding, on the other hand, can overcome stalled translation by exerting a pulling force on the nascent polypeptide chain (Goldman et al., 2015; Wruck et al., 2017). The NT^*^ domain folds very efficiently and adapts a tight conformation in the intracellular environment (Kaldmäe et al., 2020), and may therefore provide such a pulling force when fused to an inefficiently translated protein. This idea is supported by the fact that NT^*^-p53, but not p53-NT^*^, is efficiently translated *in vitro*. However, the ribosome exit tunnel can accommodate 25-30 residues of the extended nascent protein chain, which implies that co-translational folding of the NT^*^ domain alone is unlikely to be able to affect translation of the following ~90 disordered residues of p53, including the proline-rich domain at positions 61-92. To understand how this extended disordered region could be affected by the presence of an N-terminal NT^*^ domain, we therefore examined the structure of the NT^*^-p53 fusion protein. Insights from cross-linking, conformation-specific antibody studies, EM, MD simulations, and IM-MS show that NT^*^ has no marked effect on the tetrameric structure of the p53 core domains but engages in multiple interactions with the disordered N-terminus. In the fusion protein, the N-terminus wraps around the NT^*^ domain, resulting in an essentially globular protein structure. This interpretation is supported by observations from bacterial expression and computational modelling of B-Raf and K-Ras. NT^*^ significantly increases expression of B-Raf, which has a disordered N-terminal segment, and induces a globular structure in the N-terminus of the fusion protein. The K-Ras N-terminus, on the other hand, folds independently of NT^*^, resulting in efficient expression regardless of the presence of an NT^*^ tag. We therefore conclude that NT^*^ facilitates efficient translation by promoting co-translational folding through non-specific contacts with disordered N-terminal segments. The proposed mechanism would be analogous to the role of the spindle when spinning yarn, as winding of the disordered domains around the spider silk domain “spindle” moves the nascent protein chain along during translation.

Taken together, we have demonstrated how inducing co-translational folding can overcome poor translation efficiency of partially disordered proteins, opening the door to new structural investigations of particularly challenging systems. Furthermore, our observation that NT^*^ increases p53 activity, but not expression levels, in human cancer cell lines suggests additional effects of decreasing N-terminal disorder in the cellular environment. Going forward, recently developed therapeutic strategies using synthetic mRNAs may benefit from the translation-enhancing strategy reported here, as the this would reduce the amount of mRNAs that have to be delivered into cancer cells in order to elicit cell death or to induce an immune response in the case of vaccines (Kong et al., 2019; Sahin et al., 2014).

## Experimental

Full experimental procedures are given in the Supplementary Information.

## Acknowledgements

ML is supported by an Ingvar Carlsson Award from the Swedish Foundation for Strategic Research, a KI faculty-funded Career Position, a Cancerfonden Project grant (19 0480), a VR Starting Grant (2019-01961), and a KI-Cancer Blue Sky Grant. BV is supported by the European Regional Development Fund - Project ENOCH (No. CZ.02.1.01/0.0/0.0/16_ 019/0000868) and Ministry of Health, Czech Republic - DRO (MMCI, 00209805). MS is supported by DoRa plus programme funded by the European Regional Development Fund and the Republic of Estonia. DPL is supported by VR grant 2013_08807. JJ and AR acknowledge support from the Center for Innovative Medicine at Karolinska Institutet (CIMED), VR and Vinnova. GC is supported by the Olle Engqvists Stiftelse.

## Notes

https://doi.org/10.5061/dryad.34tmpg4hr

### Competing Interest Statement

The authors have declared no competing interest.

