## Supplementary material for "A “spindle and thread”-mechanism unblocks translation of N-terminally disordered proteins": Fig. S

**Contents**

Materials and Methods
Supplementary Table 1
Supplementry Figures 1-6

**Materials and Methods**

*Protein sequences.*

Histidine-6 or **-10** affinity tag

MGHHHHHH**HHHH**

Hexahistidine (H10) affinity tag

MGHHHHHHHHHH

Tobacco etch virus (TEV) cleavage (/) site

ENLYFQ/S

NT*: ACCESSION CAJ90517 with mutations D40K and K62D (bold) (major ampullate spidroin 1 precursor, partial [Euprosthenops australis])

SHTTPWTNPGLAENFMNSFMQGLSSMPGFTASQLD**K**MSTIAQSMVQSIQSLAAQGRTSPN**D**LQALN
MAFASSMAEIAASEEGGGSLSTKTSSIASAMSNAFLQTTGVVNQPFINEITQLVSMFAQAGMNDVS
A

p53: ACCESSION NP_001119584 (cellular tumor antigen p53 isoform a [Homo sapiens]); TAD1-TAD2 in bold

**MEEPQSDPSVEPPLSQETFSDLWKLLPENNVLSPLPSQAMDDLMLSPDDIEQWFTEDPGP**DEAPRM
PEAAPPVAPAPAAPTPAAPAPAPSWPLSSSVPSQKTYQGSYGFRLGFLHSGTAKSVTCTYSPALNK
MFCQLAKTCPVQLWVDSTPPPGTRVRAMAIYKQSQHMTEVVRRCPHHERCSDSDGLAPPQHLIRVE
GNLRVEYLDDRNTFRHSVVVPYEPPEVGSDCTTIHYNYMCNSSCMGGMNRRPILTIITLEDSSGNL
LGRNSFEVRVCACPGRDRRTEEENLRKKGEPHHELPPGSTKRALPNNTSSSPQPKKKPLDGEYFTL
QIRGRERFEMFRELNEALELKDAQAGKEPGGSRAHSSHLKSKKGQSTSRHKKLMFKTEGPDSD

SuperfolderGFP (sfGFP)**:**

MSKGEELFTGVVPILVELDGDVNGHKFSVRGEGEGDATNGKLTLKFICTTGKLPVPWPTLVTTLTY
GVQCFSRYPDHMKRHDFFKSAMPEGYVQERTISFKDDGTYKTRAEVKFEGDTLVNRIELKGIDFKE
DGNILGHKLEYNFNSHNVYITADKQKNGIKANFKIRHNVEDGSVQLADHYQQNTPIGDGPVLLPDN
HYLSTQSVLSKDPNEKRDHMVLLEFVTAAGITHGMDELYK

B-Raf: ACCESSION NP_004324.2 (serine/threonine-protein kinase B-raf isoform 1 [Homo sapiens]);

AALSGGGGGGAEPGQALFNGDMEPEAGAGAGAAASSAADPAIPEEVWNIKQMIKLTQEHIEALLDK
FGGEHNPPSIYLEAYEEYTSKLDALQQREQQLLESLGNGTDFSVSSSASMDTVTSSSSSSLSVLPS
SLSVFQNPTDVARSNPKSPQKPIVRVFLPNKQRTVVPARCGVTVRDSLKKALMMRGLIPECCAVYR
IQDGEKKPIGWDTDISWLTGEELHVEVLENVPLTTHNFVRKTFFTLAFCDFCRKLLFQGFRCQTCG
YKFHQRCSTEVPLMCVNYDQLDLLFVSKFFEHHPIPQEEASLAETALTSGSSPSAPASDSIGPQIL
TSPSPSKSIPIPQPFRPADEDHRNQFGQRDRSSSAPNVHINTIEPVNIDDLIRDQGFRGDGGSTTG

LSATPPASLPGSLTNVKALQKSPGPQRERKSSSSSEDRNRMKTLGRRDSSDDWEIPDGQITVGQRI
GSGSFGTVYKGKWHGDVAVKMLNVTAPTPQQLQAFKNEVGVLRKTRHVNILLFMGYSTKPQLAIVT
QWCEGSSLYHHLHIIETKFEMIKLIDIARQTAQGMDYLHAKSIIHRDLKSNNIFLHEDLTVKIGDF
GLATVKSRWSGSHQFEQLSGSILWMAPEVIRMQDKNPYSFQSDVYAFGIVLYELMTGQLPYSNINN
RDQIIFMVGRGYLSPDLSKVRSNCPKAMKRLMAECLKKKRDERPLFPQILASIELLARSLPKIHRS
ASEPSLNRAGFQTEDFSLYACASPKTPIQAGGYGAFPVH

K-Ras 4b: ACCESSION NP_004976.2 (GTPase KRas isoform b [Homo sapiens]);

MTEYKLVVVGAGGVGKSALTIQLIQNHFVDEYDPTIEDSYRKQVVIDGETCLLDILDTAGQEEYSA
MRDQYMRTGEGFLCVFAINNTKSFEDIHHYREQIKRVKDSEDVPMVLVGNKCDLPSRTVDTKQAQD
LARSYGIPFIETSAKTRQGVDDAFYTLVREIRKHKEKMSKDGKKKKKKSKTKCVIM

*Reagents*

All the reagents were purchased from Sigma-Aldrich if not stated otherwise. All constructs (Table S1) were purchased from GenScript Biotech (Netherlands) B.V.

All the protein analysis with SDS-PAGE were done by using 4–20% Mini-Protean TGX Stain-Free precast gel system (Bio-Rad, CA). The proteins were visualized using the Chemidoc Touch Imaging System (Bio-Rad, CA).

*In vitro translation (IVT)*

pETF2 and pETRC primers (Table S1) were used for cDNA amplification (Phusion High Fidelity PCR Kit, New England Biolabs, MA). 250 ng of purified cDNA (QIAquick PCR Purification Kit, Qiagen, Germany) were used for 25 µL IVT reaction based on the supplier's protocol (PURExpress In Vitro Protein Synthesis Kit, New England Biolabs, MA). The proteins were expressed for 4 h at 37 °C. 2.5 µL of the solution was mixed with reducing sample buffer, heated at 95 °C for 10 min and analysed by SDS-PAGE. For GFP fluorescence-based translation kinetics, 25 µL of IVT reactions were combined on ice into black Corning half-area 96 well plate with the clear (flat) bottom. The solution was covered with mineral oil (light oil) to avoid evaporation. GFP fluorescence kinetics were recorded for 4 h (48 cycles) at 37 °C on a Tecan Spark 20M multimode reader (Tecan Instruments, Switzerland). The excitation was conducted at 485 nm (bandwidth 20 nm), and emission was recorded at 535 nm (bandwidth 20 nm) with an integration time of 40 µs and 30 (5 × 6) flashes per well. The gain was manually set to 60%. Data were analysed with the Magellan software package (Tecan Instruments, Switzerland).

*Polysome profiling*

For polysome profiling, 50 µL of IVT reactions were incubated on a heating block at 37 ˚C for 100 min. The expression was stopped after 100 min by addition of tetracycline hydrochloride (100 µg/mL) and incubating further for 15 min. The samples were placed on ice until centrifugation. Sucrose density gradients were prepared before sample preparation on the day of the experiment using a Gradient Master 108 (Biocomp, Canada) according to manufacturer's instructions in 13.2 mL polypropylene ultracentrifuge tubes (Beckman-Coulter, CA) and stored at 4 °C. The sucrose gradients were made from 10 to 50% sucrose concentration dissolved in 20 mM HEPES pH 7.6, 100 mM KCl, 5 mM $MgCl_2$, 10 µg/mL tetracycline hydrochloride and 10 U/mL RNase inhibitor and protease inhibitor cocktail (Thermo Fischer Scientific, MA). The stopped IVT reactions were added on top of the sucrose density gradient. The gradients were centrifuged in a SW 41 swinging bucket rotor (Beckman Coulter, CA) in an Optima XPN-80 Ultracentrifuge at 36 000 rpm for 2 h at 4 ˚C with maximum acceleration and no brake option. After centrifugation, the gradients were placed at 4 °C and analysed one by one using BR-188 Density Gradient Fractionation System (Brandel Inc, MD). The parameters were 1.5 mL/min speed and 1.0 abs of resolution. The chase solution consisted of 60% sucrose. Absorbance at 245 nm was monitored using Peak Chart recording software (Brandel Inc, MD).

*Purification of NT*-TAD1-TAD2 and TAD1-TAD2*

The genes for MGH10-TEV-TAD1-TAD2 and MGH10-NT*-TEV-TAD1-TAD2 cloned into the pET26b+ expression vector were transformed into chemically competent *E. coli* BL21 (DE3) cells. Over-night cultures were inoculated 1:100 to Luria-Bertani medium containing 70 mg $L^{-1}$ kanamycin. The cultures were grown at 30 °C to an $OD_{600}$ of 0.9 before induction by 0.5 mM isopropyl-β-D-thiogalactopyranoside (IPTG) and over-night expression at 20 °C. For each construct, 500 mL culture was harvested by centrifugation at 5000 × *g* for 20 min. The pellet was resuspended in 20 mM Tris-HCl, pH 8.0 loading buffer (40 mL) and stored at -20 °C overnight. After thawing at room temperature, the cells were lysed in a cell disrupter (Constant Systems Ltd, Daventry, UK) at 30 kPsi followed by centrifugation at 27 000 × *g* for 30 min. A Ni-Sepharose (GE Healthcare, Uppsala, Sweden) column was packed from 6 mL resin and equilibrated with loading buffer. The supernatant from centrifugation was loaded on the column followed by washes with loading buffer supplemented with 10 mM imidazole. The protein was eluted (1.5 mL fractions) with loading buffer supplemented with 300 mM imidazole. Fractions with high absorbance at 280 nm were pooled and transferred to a Spectra/Por® dialysis membrane (6-8 kDa molecular weight cut-off, Spectrum Laboratories, Rancho Dominguez, CA, USA) for overnight dialysis in 5 L loading buffer at 4 °C. Samples taken from steps during expression, lysis and purification were analyzed by SDS-PAGE using 4-20% Mini-PROTEAN® TGX™ polyacrylamide gels (Bio-rad Laboratories Inc., USA) stained with Coomassie Brilliant Blue dye.

*Protein expession*

The plasmid vectors encoding p53, B-Raf, or K-Ras with or without NT* tags were transformed into BL21(DE3) competent *E. coli* (New England Biolabs, MA). Cells were grown at 30 °C, under shaking in 1 L of LB-broth supplemented with 50 µg/mL kanamycin until $OD_{600}$ reached ~0.7-0.9. Temperature was lowered to 25 °C and protein expression was induced with 1 mM isopropyl β-D-1-thiogalactopyranoside for 20 h, under shaking. Cells were harvested for 20 min at 6000 × *g*, 4 °C. The cell pellet was resuspended in 50 mM Tris pH 7.5 (30 mL of buffer per 500 mL culture) and frozen at -20 °C.

*Purification of NT*-p53 inclusion bodies (Ibs)*

The thawed cell pellets containing H6-NT*-TEV-p53 (hereinafter as NT*-p53) was resuspended in lysozyme (1 mg/mL), deoxyribonuclease I (10 µg/mL) and MgCl2 (2 mM) and incubated on ice for 1 h followed by sonication using a probe sonicator (Qsonica, CT) on ice for 5 min, 1 sec on/1 sec off at 70% power (indicator between 20−25%). After 5 min of rest on ice, the sonication was repeated followed by centrifugation for 15 min at 15 000 × *g*, 4 °C. The supernatant was discarded, and the pellet was suspended in the isolation buffer (25 mL per 250 mL culture) containing 2 M urea dissolved in 50 mM Tris, 5 mM DTT, 2% Triton X-100, 500 mM NaCl, pH 7.5. The solution was mixed and sonicated using a probe sonicator (Qsonica, CT) on ice for 30 sec in constant mode. The solution was centrifuged for 15 min at 15 000 × *g*, 4 °C. The supernatant was discarded, and the procedure was repeated once more. After 2nd centrifugation, the pellet was suspended in 50 mM Tris pH 7.5 buffer and solution transferred to the tube of known weight. The suspension was sonicated using a probe sonicator (Qsonica, CT) and centrifuged as previously. The yield was calculated by weighing the wet pellet after removal of excess buffer. From 1 L culture, approximately 2 grams of wet IBs were obtained. Isolated IBs were stored in -20 °C.

*Refolding of NT*-p53*

250 mg of IBs were dissolved in 1x denaturing buffer containing 8 M urea, 100 mM Tris, 1 mM glycine, 1 mM ethylenediaminetetraacetic acid (EDTA), 10 mM 2-mercaptoethanol, 10 mM dithiothreitol (DTT), 1 mM reduced glutathione (20 mg of IBs per 1 mL buffer). The solution was incubated at room temperature (RT) with mild continuous shaking until the pellet dissolved. The solution was centrifuged for 30 min at 48 000 × *g*, 4 °C to remove undissolved pellet and debris. The protein concentration was adjusted to ~2 mg/mL in denaturing buffer. Denatured protein was refolded by slowly switching denaturing buffer to refolding buffer as follows. On day #1, 50 mL of protein solution (2 mg/mL) in Spectra/Por dialysis membrane (6-8 kDa MWCO, Spectrum Chemicals, NJ) was placed into wide 500 mL volumetric flask, filled with 500 mL 1× denaturing buffer, and the flask was placed into 5 L beaker. Buffer exchange was carried out with by delivering buffer into the bottom of the flask while stirring using an Easy Load Masterflex L/S peristaltic pump (Cole-Parmer, IL) at a flow of approximately 0.5-0.7 mL/min. 700 mL of 0.75 × denaturing buffer was pumped through o/n at RT. On day #2, 400 mL of overflow buffer was mixed with 300 mL 2× refolding buffer containing 200 µM $ZnSO_4$, 40 mM Tris, 400 mM L-arginine, 400 mM NaCl pH 8. The obtained solution was pumped through as previously, at 4 °C. On day #3, 400 mL of overflow buffer was mixed with 300 mL 1× refolding buffer. The obtained solution was pumped through as previously. On day #4, the same was repeated as on day #3. On day #5, 400 mL overflow was mixed with 400 mL 1x refolding buffer and pumped through as previously. On day #6, 200 mL overflow was mixed with 600 mL 1× refolding buffer and pumped through as previously. On day #7, 800 mL of fresh 1× refolding buffer was pumped through. At Day #8, dialysis tube was placed into fresh 500 mL 1× refolding buffer for ~6 h. The buffer was replaced once more and dialysed o/n. On day #9, the protein solution was filtered through a 30 mm, 0.45 µm polyvinylidene difluoride (PVDF) filter (Thermo Scientific, MA), and concentrated using 100 MWCO Amicon Ultra-4 centrifugal filters (Merck-Millipore, MA) at 5000 × *g* for 30 min at 4 °C. The refolded protein (~30 mg in 5 mL) was separated from larger aggregates using the ÄKTA Start system coupled to HiLoad 26/600 Superdex 200 size-exclusion chromatography (SEC) column (GE Healthcare, IL). 1× refolding buffer was used as a mobile phase at a flow rate of 2 mL/min. To get the tetramer at the highest purity, the SEC-purification was conducted twice. Overall, ~7 mg of refolded tetrameric protein from 250 mg of IBs was obtained. All the protein quantifications were done with UV absorption

measurement (NanoDrop, Thermo Scientific, MA), applying an extinction coefficient at 280 nm of 43 025 $cm^{-1}M^{-1}$.

*Crosslinking of NT*-p53*

400 ng of refolded NT*-p53 in 10 µL of refolding buffer were crosslinked with 0.1% glutaraldehyde (1 µL of 1%) for 15 min at 37 °C. The sample was placed on ice for 10 min. 10 µL of 2x denaturing sample buffer was added to the sample and heated at 95 °C for 10 min. 10 µL (200 ng) of protein were separated by SDS-PAGE and transferred (25 V, 1.0 A, 30 min) onto 0.45 µm PVDF membrane (Bio-Rad, CA). The membrane was blocked (PBS, 5% skim milk, 0.1% Tween-20) for 1 h under gentle agitation at RT. The NT*-p53 detection was conducted with mouse anti-p53 DO-1 (in house) and polyclonal Rabbit anti-mouse IgG/HRP antibody (Agilent, DAKO).

*Mass spectrometry of NT*-p53 IBs*

NT*-p53 IBs were dissolved in 30% formic acid, 1% DTT, and diluted to a final concentration of approximately 20 µM in milliQ. Mass spectra were acquired on a Micromass LCT ToF modified for analysis of intact protein complexes (MS Vision, The Netherlands) equipped with an offline nanospray source. ESI capillaries were purchased from Thermo. The capillary voltage was 1.5 kV, the cone voltage 1.5 kV, and the RF lens 1.5 kV. The pressure in the ion source was maintained at 9.0 mbar. Spectra were visualized using MassLynx 4.1 (Waters, UK).

*Ion mobility mass spectrometry of NT*-TAD1-TAD2*

Purified NT*-TAD1-TAD2 and beta-lactoglobulin (Sigma, MO) as calibrant were buffer-exchanged into 100 mM ammonium acetate, pH 7.0, using BioSpin6 columns (BioRad, CA). IM-MS spectra were acquired on a Waters Synapt G2 travelling wave ion mobility mass spectrometer (Waters, UK) equipped with an offline nanospray source. The capillary voltage was 1.5 kV, the source presure was 3.4 mbar, and the source temperature was 80 °C. The collision energy in the ion trap was 10 V. Wave height and wave velocity were 35 V and 700 m/s in the IMS cell and 248 m/s and 10 V in the transfer. IMS gas was nitrogen with a flow of 50 mL/min. Mass spectra were visualized using MassLynx 4.1. CCS calculations were performed based on the beta-lactoglobulin monomer and dimer using PULSAR.

*Immunoprecipitation of refolded NT*-p53*

Dynabeads Protein G (Thermo Fischer Scientific, MA) (5 µL per sample) were washed three times with phosphate-buffered saline (PBS) supplemented with 0.1% Tween-20 and 0.1% bovine serum albumin (BSA). Beads were incubated with 1 µg of anti-p53 monoclonal antibodies (DO-1, PAb1620, PAb240, PAb421; in house) in 30 µL of 3% BSA for 1 h at RT on a tube rotator. Anti-HA monoclonal antibody (Sigma, H3663) was included as a negative, and sample without antibody as a double negative control. After incubation, the beads were washed as previously. 10 µL of 5 µM protein sample in the refolding buffer was mixed with the beads and incubated o/n at 4 °C on a tube rotator. Beads were washed six times as previously, followed by two times with PBS. The beads were mixed with 20 µL of reducing sample buffer and heated at 95 °C for 10 min. Magnetic beads were collected and 5 µL of supernatant was used for SDS-PAGE. The proteins were transferred (25 V, 1.0 A, 10 min) onto 0.2 µm nitrocellulose membrane (Bio-Rad). The membrane was blocked (PBS, 5% skim milk, 0.1% Tween-20) for 1 h under gentle agitation at RT. The NT*-p53 detection was conducted with DO-1 HRP antibody (Santa Cruz Biotechnology, sc-126).

*Preparation and TEM imaging of NT*-p53 tetramer.*

Refolded NT*-p53 tetramer was injected into Superose 6 column (GE healthcare, IL) connected to the ÄKTA Pure system (GE healthcare, IL), and NT*-p53 was eluted out with 20 mM ammonium acetate pH 8.0. To get homogenous NT*-p53 tetramer species, a narrow peak fraction (~ 100 µL) was collected with a concentration of around 30 µg/mL. For negative staining grid preparations, 4 µL homogenous NT*-p53 tetramer were applied onto glow-discharged continuous carbon-coated copper grids (400 mesh, Analytical Standards) and allowed to absorb for 2 min. The grids were subsequently blotted with pre-wetted filter paper and washed with two drops of milli-Q water sequentially. After blotting away the washing drops, the grids were negatively stained with one drop of 2% (w/v) uranyl acetate for 45 sec, subjected to final blotting and air-drying. The sample was imaged using a Jeol JEM2100F field emission gun transmission electron microscope (Jeol, Japan) operating at 200 kV. A total of 30 micrographs were recorded using a Tietz 4k × 4k CCD camera, TVIPS (Tietz Video and Image Processing Systems, Germany) at a magnification of 85 000 (1.76 Å per pixel) and 0.8 – 2.0 µm defocus.

*Processing of single particle images*

All recorded micrographs were imported to EMAN2 (version 2.3) for single particle analysis [1]. After estimating defocus with e2evalimage.py, imported micrographs were subjected to particle picking, during which single particles in different orientations were manually selected from the images using e2boxer.py (4733 particles). The contrast transfer function (CTF) correction and measuring of additional parameters were performed on particle data (boxed out region containing particles, 180×180 pixels) using e2ctf.auto.py. Reference-free class averages (2D refinement) based on the selected 6519 CTF-corrected particles (low-pass filtered to 20 Å) was performed using both e2refine2d_bispec.py and e2refine2d.py. Some 2D classes show an approximate 2-fold symmetry, in agreement with previous study of p53 and the biochemical data. 2D classes generated by e2refine2d_bispec.py were used as the input for building the 3D initial model using e2initialmodel_sgd.py with D2 symmetry. Two rounds of 3D refinement were performed using e2refine_easy.py applying D2 symmetry aiming at a final resolution of 20 Å. The first round of 3D refinements was performed with input data low-pass filtered to 12 Å (pixel size 3.77 Å), while the second round of refinement was carried out with data low-pass filtered to 7 Å (pixel size 2.475 Å). The final map from the first round of refinements was used as starting model in the second round. The resolution was determined based on a Fourier Shell Correlation (FSC) value of 0.143 [2], following the gold standard FSC procedure implemented in EMAN2 [3]. Structures of the p53 core tetramer (PDB ID 3KMD) and the NT domain (PDB ID 2LPJ) were fitted into the density using the “Volume viewer” and “Fit in map” tools in UCSF Chimera V1.11.2.

*Construction of the NT*-TAD1-TAD2-PRD, NT*-B-Raf$_{1-40}$ and NT*-K-Ras$_{1-40}$ models*

The NMR structure of spider silk N-terminal domain (NT; PDB ID: 2LPJ ) was downloaded from the Protein Data Bank. The lowest energy state structure (Model 1) from the reported ensemble of twenty NT structures was selected and two in-silico mutations “D40K and K65D” were incorporated (NT*) as per the recombinant protein used in the current work. A 97 amino acid sequence comprising of TEV cleavage site ($^{1}$ENLYFQS$^{7}$) along with the transactivation domains (TAD1 and TAD2), and the proline rich domain (PRD) of p53 was considered for modelling NT*-TAD1-TAD2-PRD structure. Extended state conformations of “TEV+TAD1 (residues 1-40)”, TAD2 (residues 41-70) and PRD (residues 71-90) polypeptide segments were generated separately with the LEaP module of AMBER18 [4]. The MDM2 binding site (18TFSDLWKLL26) located in TAD1 was assigned a helical backbone conformation. Modelling of NT* and the N-terminal domains from p53 was done in a sequential manner (Figure S6). TEV+TAD1 was first attached via a peptide bond with NT* and subjected to explicit solvent MD simulations (details mentioned below) to generate NT*-TAD1 structures. An end-state representative structure of this system was then attached to TAD2 and simulated to obtain “NT*-TAD1-TAD2” structure. PRD was finally added to the construct to model “NT*-TAD1-TAD2-PRD” structure. A 47 amino acid sequence comprising of TEV cleavage site ($^{1}$ENLYFQS$^{7}$) and the first 40 residues from the N-terminus of B-Raf and K-Ras was used to model the NT*-B-Raf$_{1-40}$ and NT*-K-Ras$_{1-40}$ structures repsectively.These models were generated by sequentially substituting the residues from B-Raf and K-Ras sequence onto the backbone scaffold of NT*-TAD1 structure.

*Molecular dynamics simulations*

The N- and C-terminus of “NT*-TAD1”, “NT*-TAD1-TAD2”, “NT*-TAD1-TAD2-PRD”, NT*-B-Raf$_{1-40}$ and NT*-K-Ras$_{1-40}$ models were all capped with ACE (acetyl) and NME (N-methyl) functional groups respectively. These models were placed in the centre of an octahedral box whose dimensions were fixed by maintaining a minimum distance of 8 Å between any protein atom and the box boundaries. TIP3P water model [5] was used to solvate the structure. The net charge of the systems were -12e (NT*-TAD1), -21e (NT*-TAD1-TAD2 and NT*-TAD1-TAD2-PRD), -11e (NT*-B-Raf$_{1-40}$) and -10e (NT*-K-Ras$_{1-40}$), which were neutralized by adding 12 Na+, 21 Na+, 11 Na+ and 10Na+ counter ions respectively. All-atom molecular dynamics simulations were carried out using the PMEMD module through the AMBER18 suite of programs employing ff14SB force field parameters [6]. The systems were energy minimized using steepest descent followed by conjugate gradient schemes, heated to 300 K over 30 ps under NVT ensemble and equilibrated for 200 ps in NPT ensemble. The final production dynamics was run for 1 μs each under NPT conditions. NT*-TAD1, NT*-B-Raf$_{1-40}$ and NT*-K-Ras$_{1-40}$ systems were all simulated in triplicates with different initial velocities. Simulation temperature was maintained at 300 K using Langevin dynamics [7] (collision frequency: 1.0 ps$^{-1}$), and the pressure was kept at 1 atm using weak-coupling [8] (relaxation time: 1ps). Periodic boundary conditions were applied on the box edges and Particle Mesh Ewald (PME) method [9] was used to compute long-range electrostatic interactions. SHAKE algorithm [10] was used to constrain all the bonds involving hydrogen atoms and the equation of motion was solved with an integration time step of 2fs.

*Mammalian cell lines and transfection*

NCI-H1299 human non-small cell lung carcinoma cells, kindly provided by the p53 Laboratory at the Agency for Science, Technology and Research, Singapore, were cultured and maintained in DMEM high glucose, supplemented with 10% fetal calf serum (Hyclone, GE healthcare, UK), 2mM L-glutamine, 100 IU penicillin and 100 μg/mL streptomycin at 37°C 5% $CO_2$. Transfection of described constructs (Table S1) was performed at 37°C 5% $CO_2$ by use of Lipofectamine 3000 (Thermo Fischer Scientific, MA) according to instructions provided by the manufacturer upon reaching 70-90% cell confluency. H1299 cells grown on 8-well Permanox slides (Thermo Fischer Scientific, MA) were transfected with 100 ng/well (pCMV-GFP without insert) or 200 ng/well (all other constructs) plasmid DNA in Gibco OptiMEM (31985-047, Thermo Fischer Scientific, MA) for 4 hours before medium was changed to DMEM without phenol red.

*Immunocytochemistry and immunofluorescence imaging*

Cells were fixed 8 hours post transfection with 4% paraformaldehyde (room temperature, 15 min) and permeabilized with 0,1% Triton X-100 in 5% BSA PBS solution (room temperature, shaking at 20 rpm, 10 min), followed by subsequent blocking with 5% bovine serum albumin in PBS containing 0,1% polysorbate 20. Used primary antibodies were mouse anti-p53 [DO-1] (ab1101, Abcam, MA, 1:100) and rabbit anti-TIMM23 (11123-1-AP, Proteintech, IL, 1:50), and were applied in PBS with 0,1% polysorbate 20 and 1% BSA, and left for overnight incubation at 4°C. Secondary labeling was performed for detection with Alexa 488- and Alexa 647-conjugated goat anti-mouse or anti-rabbit IgG (Life Technologies, 1:1000, room temperature, 1 hour). Cells are consecutively stained with DAPI nuclear staining (D1306, Life Technologies, 1:10000) and coverslips were mounted with ProLong Gold (Thermo Fischer Scientific, MA). Three washes were performed after each labeling step with PBS containing 0,1% polysorbate 20. Control experiments for each antibody staining were performed with exclusively secondary antibodies. Confocal images were acquired though immunofluorescence microscopy using a Nikon Eclipse Ti series inverted microscope (Nikon), equipped with Crest X-light V2 series confocal unit (Nikon). Images were acquired using an S Plan ELWD 60X/0,70 objective (Nikon) and a Zyla sCMOS camera (Andor). NIS-Elements Advanced Research 4.60.00 64-bit software (Nikon) was used for image analysis.

*Luciferase reporter assay*

H1299 cells were maintained in Dulbecco's modified Eagle's medium (DMEM) with 10% (v/v) foetal calf serum (FCS) and 1% (v/v) penicillin/streptomycin. The cells were seeded at

1.0 × $10^5$ cells/well in 6-well plates, 24 h prior to transfection. Cells were co-transfected with the relevant p53 plasmids, LacZ reporter plasmid and luciferase plasmid using Lipofectamine 2000 (Thermo Fisher Scientific, MA) according to the manufacturer’s instructions. The cells were harvested 24 h after transfection and β-galactosidase activities were assessed using the Dual-light System (Applied Biosystems, CA) according to the manufacturer's protocol. The β-galactosidase activity was normalized with luciferase activity for each sample. To check for expression levels of relevant proteins via western blot, 5 µg of the cell lysates were probed for p53 with horseradish peroxidise conjugated DO1 antibody (Santa Cruz, TX) and β-actin-peroxidase antibody (Sigma, MO).

**Table S1.** DNA constructs used in this study

| Use | Construct | Vector | Restriction sites |
|---|---|---|---|
| **IVT**† | H6-NT*-p53-sfGFP<br>H6-p53-sfGFP<br>H6-NT*-sfGFP<br>H6-NT*-p53<br>p53-NT*-H6 | pET26b(+) | *Nde*I, *BamH*I |
| **Expression in BL21** | H6-NT*-TEV-p53<br>H10-NT*-TAD1-TAD2<br>H10-TAD1-TAD2<br>H6-p53<br>H6-B-Raf<br>H6-NT*-B-Raf<br>H6-K-Ras<br>H6-NT*-K-Ras | pET26b(+) | *Nde*I, *BamH*I |
| **Expression in human cell lines** | H6-NT*-TEV-p53<br>H6-p53$^{R273H}$<br>H6-p53 | pCMV6-A-GFP‡ | *Sgf*I, *Rsr* |

†For cDNA amplification (used in IVT restriction) the following primers were used:

pETF2 5´CATCGGTGATGTCGGCGAT´3

pETRC 5´CGGATATAGTTCCTCCTTTCAGCA´3

‡Insert under CMV promoter; turboGFP under separate (SV40) promoter used as selection marker.

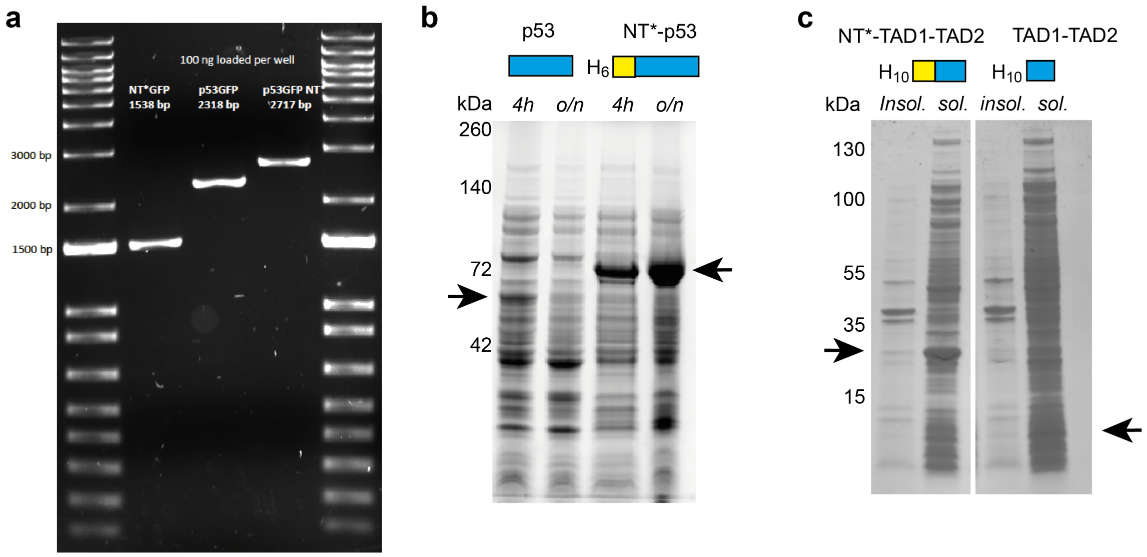


**Figure S1. IVT input and bacterial expression of p53 NT*-p53, TAD1-TAD2, and NT*-TAD1-TAD2.** (a) Agarose gel showing the PCR products used as input for the IVT reactions. (b) Expression of p53 and NT*-p53 in BL21 cells under identical conditions show a minor amount of p53 after 4 h expression, which is degraded after 24 h. NT*-p53, on the other hand, is expressed rapidly and accumulates without notable degradation. (c) SDS PAGE analysis of the soluble and insoluble fractions of *E. coli* expressing either $His_{10}$-TAD1-TAD2 or $His_{10}$-NT*-TAD1-TAD2 shows a visible band for the NT*-tagged construct. Arrows indicate the expected molecular weight of each protein. The yields of $His_{10}$-TAD1-TAD2 and $His_{10}$-NT*-TAD1-TAD2 from 0.5 L of bacterial culture were 4 and 23 mg, respectively.

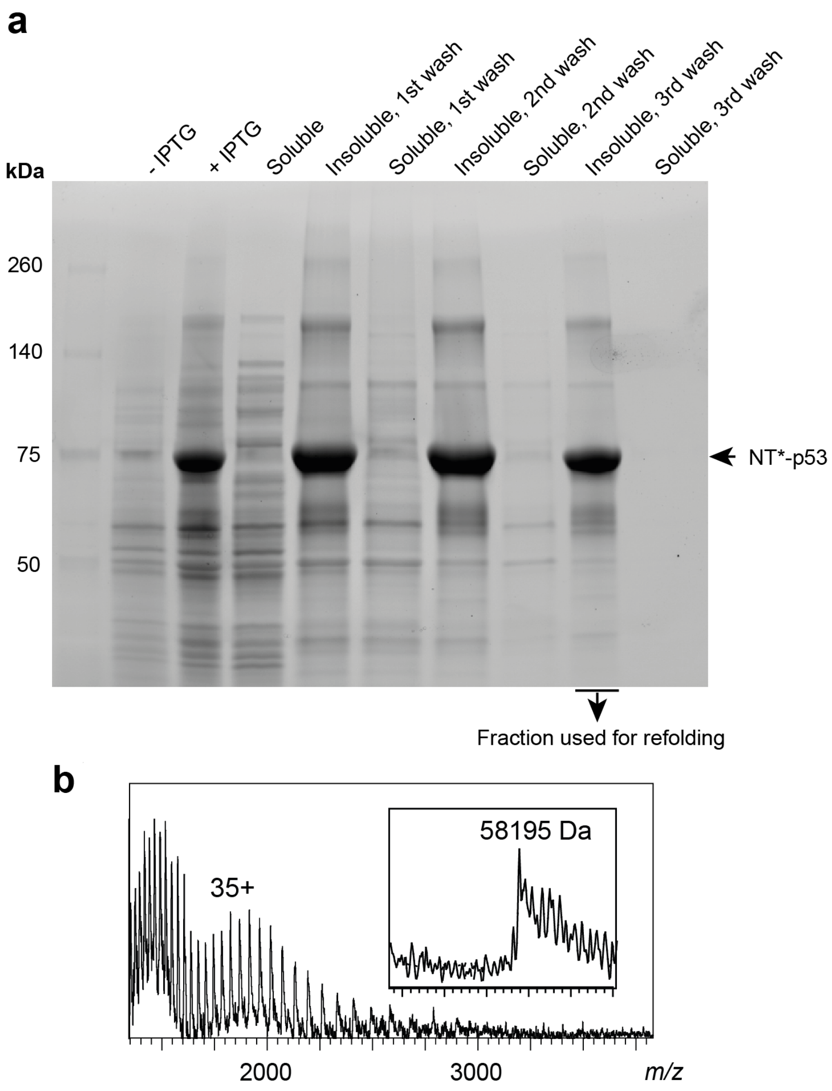


**Figure S2. Purification of NT*-p53 from inclusion bodies.** (a) SDS-PAGE showing total lysates of BL21(DE1) cells before and after induction of NT*-p53 expression. Upon expression, the protein (arrow) is found exclusively in the insoluble fraction. Multiple rounds of sonication and washing yield >90% pure NT*-p53 in the insoluble fraction which was then used for refolding. (b) Mass spectrometry of the insoluble fraction reveals pure, full-length NT*-p53 protein.

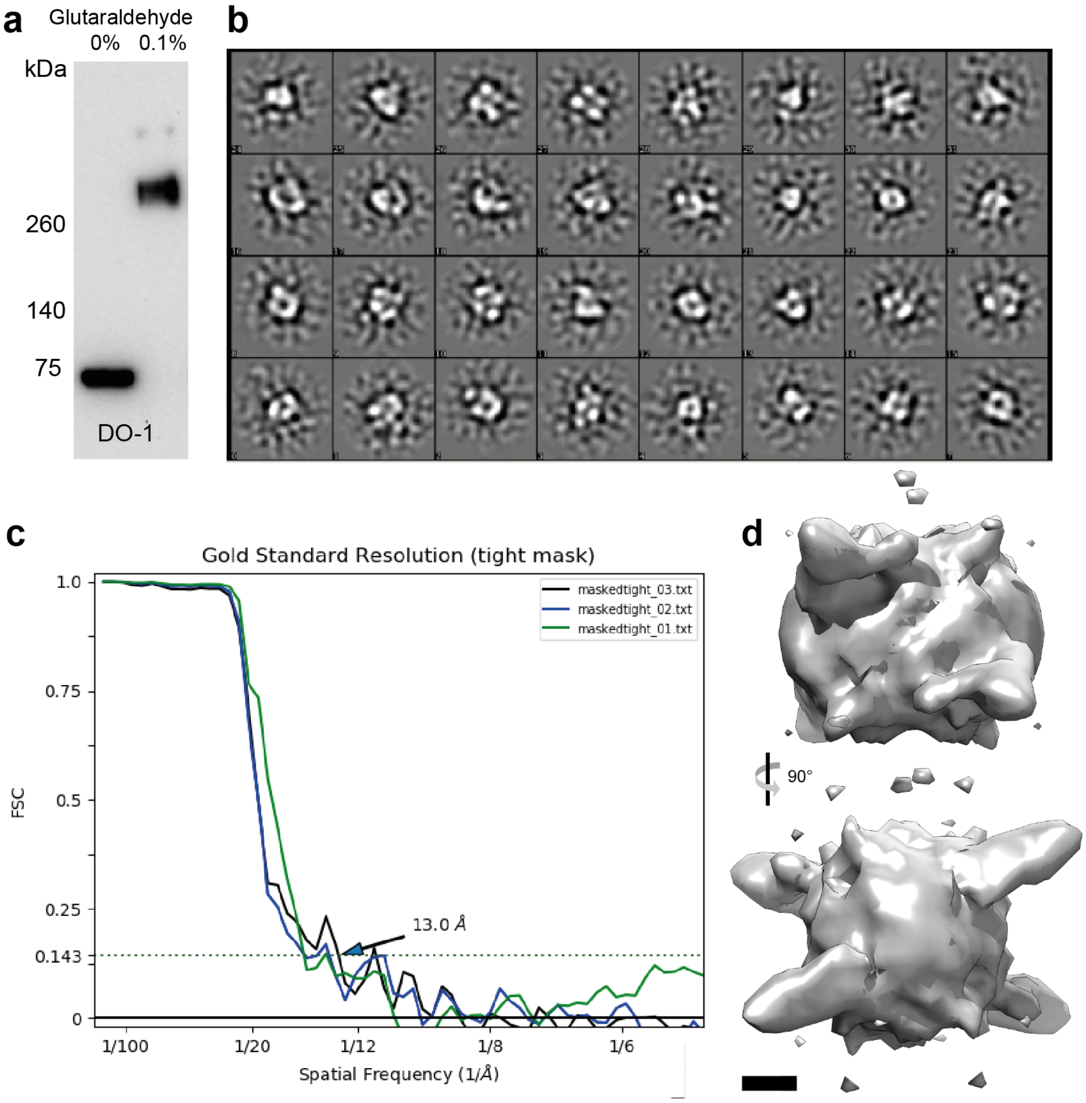


**Figure S3. Negative stain EM of NT*-p53. (**a) Western blot of refolded NT*-p53 before and after crosslinking with glutaraldehyde confirms tetramer formation. (b) Class averages of NT*-p53 observed by uranyl acetate staining of refolded NT*-p53. (c) The Fourier shell correlation (FSC) indicates the resolution of ~13 Å for the reconstructed 3D density map. The determined resolution was calculated from the curve at FSC = 0.143 (dotted line). (d) 3D density map of NT*-p53 tetramer with dihedral (D2) symmetry. The map was based on 6519 particles extracted from images recorded on a CCD camera. The scale bar (black) is 20 Å.

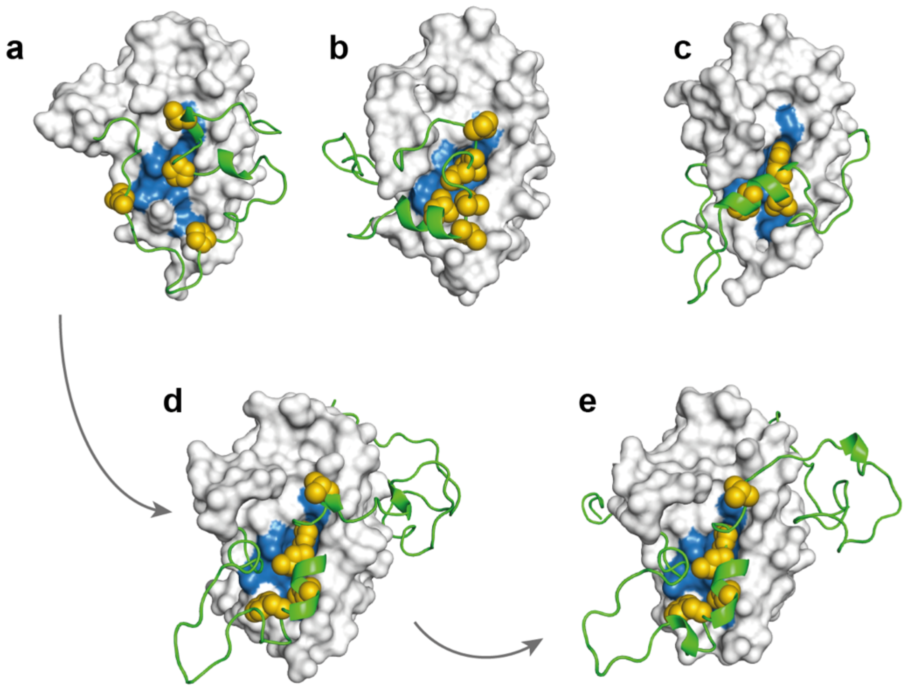


**Figure S4. Modelled structures of NT* and p53 N-terminal domains.** End state representative states from MD simulations of (a, b and c) NT*-TAD1 performed in triplicates, (d) NT*-TAD1-TAD2 and (e) NT*-TAD1-TAD2-PRD structures. The sequential manner of modeling the structure of NT* and the different domains from a representative end state "NT*-TAD1" structure is highlighted with arrows. NT* and the p53 N-terminal domains along with the TEV sequence are shown in surface and cartoon representations respectively. Hydrophobic residues in TAD1 domains are shown as yellow spheres. The hydrophobic surafce patch formed by residues M41, I44, M48, I52, L55, L66, A68, L69, A72, F73 and M77 on the dimerization interface of NT* is highlighted in blue.

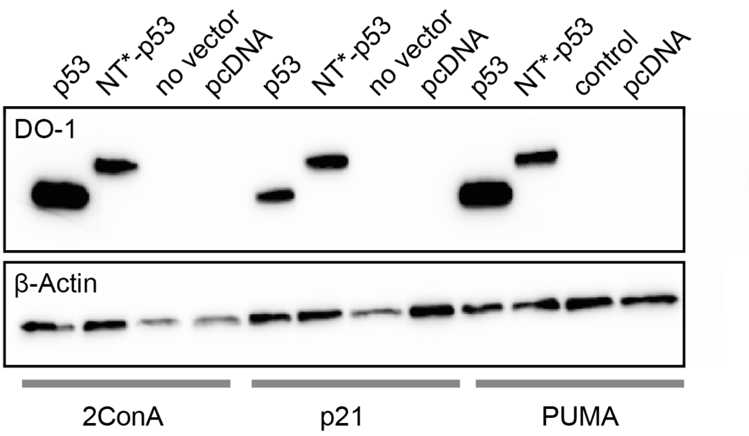


**Figure S5. NT* does not increase p53 levels in H1299 cells.** Western blot of 5 μg cell lysate showing p53 expression in H1299 reporter cells 24 h after transfection with either wt p53 or NT*-p53.

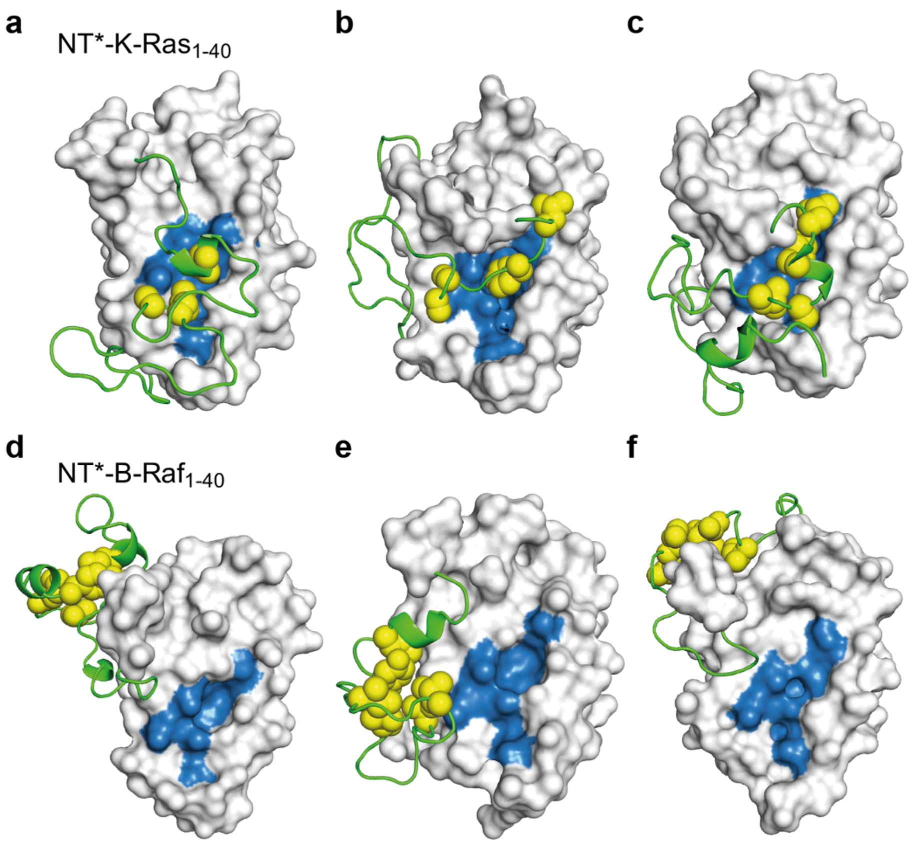


**Figure S6. Modelled structures of NT* with the N-terminal residues of B-Raf and K-Ras.** End state representative states from three independent MD simulations of (a, b and c) NT*-B-Raf$_{1\text{-}40}$ and (d, e and f) NT*-K-Ras$_{1\text{-}40}$ structures. NT* and the N-terminal residues from B-Raf and K-Ras along with the TEV sequence are shown in surface and cartoon representations respectively. The Hydrophobic residues in TEV+B-Raf and TEV+K-Ras are shown as yellow spheres. The hydrophobic surface patch formed by residues M41, I44, M48, I52, L55, L66, A68, L69, A72, F73 and M77 on the dimerization interface of NT* is highlighted in blue.
